# Forest characteristics predict tri-colored bat activity within novel Colorado habitats

**DOI:** 10.1101/2023.05.11.540259

**Authors:** Amanda J. Bevan Zientek, Alexandria B. Colpitts, Rick. A. Adams

## Abstract

Climate change and other anthropogenic pressures are altering species distributions. Several studies have indicated that tri-colored bats (*Perimyotis subflavus*) are extending their distributional range westward in the United States. Montane and subalpine habitats consist of a mosaic of forest types including lodgepole pine woodlands and montane meadows, which provide an opportunity to study how a newly arriving species, typically associated with lowland riparian systems, is adapting to novel environmental conditions. The objective of this study is to document tri-colored bat activity in these novel habitats using acoustic surveys and to quantify what factors are influencing activity patterns in habitats and at elevations not previously documented. We selected sites at 2700m elevation that differed in stand structure, driven primarily by beetle kill outbreaks, and were in various stages of secondary succession. We used acoustic monitoring to model habitat activity patterns using nonparametric multiplicative regression. Results showed that tri-colored bats used meadows and lodgepole pine stands undergoing secondary succession following bark beetle outbreaks. Activity was highest in meadows and early time-since-kill (TSK) forests in the beginning of the survey period, and tri-colored bats had increased activity in late TSK forest habitats at the end of the survey period in early August. Temporally, activity was lowest during the middle of the survey period (mid-July) indicating that tri-colored bats moved away from our study area. However, activity significantly returned by the end of the survey. Our study demonstrates that tri-colored bats are not restricting their activity to lower elevation riparian areas in the Colorado foothills, but instead appear to be using high elevation habitat types in areas dominated by lodgepole pine and subalpine meadows. We hope this study will support conservation efforts for this species following the proposed U.S. Fish and Wildlife Service listing.

## INTRODUCTION

Human-driven landscape changes and climate warming have increased western leading-edge movements of angiosperms (Fei et al. 2017) resulting in expansive forest and woodland corridors along rivers and streams in the Great Plains of North America. This expansion is associated with westward range extensions of many woodland animals, including mammals (see Benedict et al. 2000 for review). Indeed, species distributions have been changing worldwide (Jetz et al. 2007, Visconti et al. 2016, Pacifici et al. 2020), and bats are no exception (Rebelo et al. 2010, Razgour et al. 2013, Sherwin et al. 2013, Ancillotto et al. 2016).

Tri-colored bats (*Perimyotis subflavus*), a small foliage-roosting species historically distributed throughout eastern and midwestern North American deciduous forests (Fujita and Kunz 1984), have been found as isolated records in several western states including New Mexico, Texas, South Dakota, and Colorado (Geluso et al. 2005, Valdez et al. 2009, Fitzgerald et al. 1989, Armstrong, Adams, & Taylor 2006, Adams et al. 2018). Westward range movements appear to occur via riparian corridors along major river systems (Geluso et al. 2005, Kurta et al. 2007). Along the westward front, invasions into previously unexperienced and therefore novel habitats, as well as climates, are likely to ensue. Although in many cases, the eastern plains of the Rocky Mountain states provide rich riparian corridors along streams, rivers, and ponds, as one approaches the eastern foothills of the Rocky Mountains there ensues a dramatic change in altitude, habitat, and ecotone compression that in many cases acts as a significant dispersal barrier for many species, even large birds (Machado et al. 2018).

Tri-colored bats show signs of establishing reproductive populations along riparian systems in eastern Colorado. A female tri-colored bat with twins was found roosting in a deciduous tree in an urban riparian setting in Boulder County, Colorado, USA (Adams et al. 2018). The first record of a tri-colored bat in Colorado occurred in 1987 in Weld County (Fitzgerald et al. 1989), and deceased individuals were found in 2004 and in 2017 in riparian areas in Boulder County, CO (Armstrong et al. 2006, Adams et al. 2018). Since 1995, annual bat surveys have been conducted at lower elevation sites (1645m - 2130m) in the foothills of Boulder County consisting of ponderosa pine woodland and mountain riparian habitats with the first acoustic detection of PESU occurring in 2012 (Adams, unpl. data). Following the discovery of a reproductive female in 2017 along Boulder Creek, a follow up acoustic survey recorded 707 tri-colored passes over 10 nights in August which was a notable increase. In September, at the same site, a foraging nonreproductive female was captured in a mist net. The following month, a male tri-colored bat was found approximately 35 km north of the Boulder Creek site (Adams et al. 2018). Subsequent acoustic surveys have indicated an increase and continued presence of tri-colored bats in foothills habitats (1980–2400m) since 2017 (Rick A. Adams, [University of Northern Colorado, Greeley, CO], personal observation). Additional acoustic surveys in 2019 detected tri-colored bat movement into higher elevation habitats (2700m) including the use of lodgepole pine forests recovering from a severe bark beetle disturbance event. In 2020, we began to survey for tri-colored bats in these higher elevation sites including montane and subalpine habitats of Larimer County, CO that had been decimated by bark beetles. Although tri-colored bat activity indicates their continued presence within their expanded range, habitat associations for this species within the mountain west are thus far unstudied until now.

Tri-colored bats were once an abundant species throughout the midwestern and eastern United States, but populations have experienced precipitous declines up to 95% where white-nose syndrome (WNS) occurs (Powers et al. 2015, O’Keefe et al. 2019). Recent documentation of reproduction of this species in Colorado, which has not yet suffered from WNS population losses, indicates these habitats may provide refuge and an opportunity for population recovery. Although initial detections of tri-colored bats were in eastern Colorado along riparian creek and river systems (Adams et al. 2018), acoustic surveys indicate their expansion into novel habitats at elevations not previously occupied. Declining species are expected to have sparse distribution patterns due to population retractions to refuge areas least impacted by threats or to optimal habitats (Channell and Lomolino 2000, Wilson et al. 2004). Additionally, species occupying the periphery of their geographical range tend to have high spatio-temporal variability potentially due to poorer habitat quality (Brown et al. 1984, Brown et al. 1995, Channell and Lomolino 2000). As we suspect that tri-colored bats are at the edge of their geographic range and have experienced precipitous declines from WNS, we might expect that activity rates would be highly variable and this species might undergo adaptations to novel niche spaces in the Southern Rocky Mountain foothill and montane habitats (Philips et al. 2010, Miller et al. 2020). It is unclear how stable tri-colored bat activity is within their expanded range and how variable habitat use patterns are and the associated habitat characteristics tri-colored bats tolerate compared with those found within their historical range.

The objective of this study is to document the presence/absence of tri-colored bats at higher elevations consisting of montane and subalpine forests that would predictably be novel habitats for this species and to determine what habitat features predict tri-colored bat activity. Specifically, we hypothesize that tri-colored bat populations will not be persistently active during the active season (late spring to early fall) at these high elevation sites. Alternatively, if tri-colored bats are persistently active at our high elevation sites throughout the active season, they will show habitat-specific activity patterns.

To test these hypotheses, we conducted acoustic surveys in various habitat types with multiple detectors gathering data simultaneously and tested the following predictions: 1) mean passes per night recorded for each site will be significantly different across independent sampling periods and 2) habitat features associated with tri-colored bat activity will vary across the active season.

## MATERIALS AND METHODS

### Study area and design

Our field sites were located in the Red Feather Lakes Area of Roosevelt National Forest, Larimer County, Colorado, USA between 2,705–2,827 meters in elevation (Fig. 1). Sites are dominated by lodgepole pine forests (*Pinus contorta*) that have been heavily affected by mountain pine beetle (*Dendroctonous ponderosae*) outbreaks resulting in a mosaic of secondary successional habitats. This mosaic allowed for quantifying how newly founded tri-colored bats are adapting to elevations and habitats not previously inhabited by this species.

Using the U.S. Forest Service’s Aerial Vegetation Survey dataset (Johnson & Wittwer 2006), we identified 1-ha study sites that were severely affected by bark beetles (≥ 50% stand mortality) from 2008 – 2012 and represented successional lodgepole pine recovery stages (Fig.1). We categorized habitat types as “early time-since-kill (TSK)”, “mid TSK”, and “late TSK” successional stages using a combination of time-since-kill (quantified using the methodology described below) and visual estimates of the degree of understory response. To fill-in a continuum of secondary successional phases, we also surveyed subalpine meadows that were dispersed throughout the study area. Ground surveys confirmed sites were severely affected by bark beetles and not by fire by evaluating that overstory trees and large fallen snags had pitch tubes present but lacked scorch marks.

### Habitat characteristics

We analyzed 0.28 ha plots at the center of each one-hectare area for elevation, slope, and aspect. We quantified the degree of canopy closure using a spherical crown densiometer and averaged across measurements taken in the cardinal directions. To estimate degree of openness, the distance between “nearest neighbor” trees was measured in approximately the four cardinal directions from the plot center and then was averaged as a measure of flight corridor width. Transformed aspect was calculated by sin(aspect in degrees/(360°/π)). This converts aspect to an index where south-facing slopes receive a high score, and north-facing slopes receive a low score (Hayes & Adams 2015). Incident radiation was calculated using aspect, latitude, and slope in radians and was then transformed from the logarithmic to the arithmetic form (McCune & Keon 2002). We collected these characteristics to include in our models to ensure habitat characteristics and not environmental characteristics are influencing tri-colored bat activity.

Vegetations surveys were conducted along a 30 x 1 m belt transect that was centered on each one-hectare plot, and transect directions were chosen using a random number generator (1- 360). Along the belt transect the diameter at breast height (DBH) of each standing tree (alive and dead) was recorded. In addition to DBH, the height, state (alive or dead) and, if beetle killed, the time since kill were recorded. The time since kill (TSK) was quantified by estimating the percentage of remaining needles, twigs, and branches and degree of faded needles in the crown and recorded (Klutsch et al. 2009). Average percent beetle kill, average TSK, maximum TSK, mode TSK, basal area, and average overstory height were then calculated for each transect and averaged. The percentage cover of shrubs, saplings/seedlings, forbs, grasses, bare ground/rock, moss and lichens, medium woody debris (large branches and woody materials less than ≤ 0.91 m long), and fine woody debris (small twigs and branches) were estimated at alternating one-meter intervals. The height of shrubs, forbs, and saplings/seedlings were also recorded for cover ≥ 5.0 cm tall. Percent cover estimates and heights were average for each class for each transect and then averaged between the two transects. The volume of coarse woody debris (CWD) was also quantified along belt transects, which included any down and dead wood debris that were ≤ 7.62 cm in diameter and ≥ .9144 m long for decay classes one through four and for decay class five and ≥12.7 cm above the duff layer with a length ≥ 1.5 m (Woodall & Monleon 2007). The diameter of each piece at the point of intersect with the transect was recorded and averages were calculated for the total volume of coarse woody debris and for each decay class (Ståhl et al. 2001). These characteristics are standard forestry measurements collected to study which features might influence bat habitat use and see Supplementary Data SD1 for these data.

### Acoustic surveys

In 2020, we used three Wildlife Acoustics SM2BAT detectors (Wildlife Acoustics, Maynard, Massachusetts) fitted with SMX-US microphones to record bat activity from June 1^st^ to August 14^th^ at 12 sites (4 early TSK, 3 mid TSK, 3 late TSK, and 2 meadow sites). No sampling occurred after the evening of August 14^th^ due to the Cameron Peak Fire which burned over 84,400 hectares and is Colorado’s largest fire in history. One detector was placed at the center of each of the three one-hectare plots to record simultaneously until detectors (3) were relocated or unless the batteries died before the end of the sampling period. Sites were surveyed a maximum of three times (N = 10 total sampling periods). However, sampling period length did vary at the beginning of the active season to account for poor weather conditions. Following the second sampling period, detectors recorded for seven nights before being relocated. The 10^th^ and final sampling period consisted of only five nights due to mandatory Cameron Peak wildfire closures. The sampling order of numbered sites was chosen using a random number generator. Bat echolocation passes were identified to species using Sonobat 4.4 (Szewczak and Szewczak 2018) with a 90% call matching threshold. Visual confirmation of a subset of passes was also done using procedures outlined in Reichert et al. (2018), call characteristics as described by Szewczak et al. (2011), and recordings from Adams et al. (2018, Supplemental Fig. 1).

## Quantitative analyses

### Spatial and temporal patterns

Maps were made using a combination of QGIS 3.22 (QGIS Development Team 2022) and Google Earth Pro (Google Earth Pro 7.3 2019). To display the mean number of passes detected per night for a given sampling period, a point shapefile with detector locations was created on top of an imported Google Map Satellite base map. Mean passes per night for the map’s respective sampling period was included with each point on the shapefile, each of which represent a site at which an acoustic detector was placed. The number of passes at each site were visually displayed by selecting a graduated color symbology type weighted by the mean passes per night data.

Total number of tri-colored bat passes recorded each night were summed with mean passes per night (PPN) and the associated standard deviation calculated by dividing the total passes recorded by the number of survey nights. To address Prediction 1, we conducted a Kruskal-Wallis test on mean PPN detected across each sampling period for sites that were surveyed three times and a U Mann Whitney test for sites that were surveyed twice in R (R Studio Version 4.0.2) to test for significant differences.

### Habitat features used for predictive modelling of bat activity

We ran a nonparametric multiplicative regression (NPMR; HyperNiche 2.30; McCune and Mefford 2009) which uses a local multiplicative smoothing function with a leave-one-out cross-validation to estimate tri-colored bat nightly detections and to test Prediction 2 (Berryman and McCune 2006).

HyperNiche also screens for highly correlated predictors during the leave-one-out approach while determining model fit. We used a Gaussian weighting function with a local mean estimator to identify forest characteristics that might predict our response variable and then expressed fit as a cross-validated R^2^ (xR^2^). Local mean distance measures are appropriate for response data as they do not generate estimates outside of the range of observations and will not produce less than zero abundance predictions (McCune 2006). The cross-validated R^2^ is calculated by excluding each data point from the basis for the estimate of the response at that point. NPMR fits tolerances, rather than coefficients in a fixed equation, which are the standard deviations used in the Gaussian smoothers (Berryman and McCune 2006). Tolerances indicate how broadly information can be used from nearby areas in predictor space. For example, high tolerances indicate information is used from a large neighborhood of data points and small tolerances indicate narrow neighborhood sizes (McCune 2006). A predictor was included in the final model if it increased xR^2^ by at least 0.03 (Averett et al. 2016), and predictor importance was evaluated by its sensitivity which is calculated by the ratio of the relative mean difference in the response to the relative mean difference in the predictor. A predictor with a sensitivity of one indicates that we would expect a 1:1 change in the response variable with changes in the predictor variable. For example, a 10% change in a predictor variable with a sensitivity of one would lead to a 10% change in the response variable, whereas a sensitivity of zero indicates that that predictor has no effect on the model (McCune 2006). Finally, we produced 3-dimensional response contour graphs for increased interpretability. We considered decreasing neighborhood sizes for generating response surfaces (and not during the model fitting phase) to reflect increased species tolerances to improve interpretability of 3-dimensional surfaces where appropriate but restricted this reduction to a minimum N* of 0.5 (McCune 2006).

## RESULTS

A total of 439 tri-colored bat passes were auto identified by Sonobat. To further validate passes, we selected a random sample of five percent of passes over the span of the survey period (22 independently detected passes) to compare with acoustic recordings from Boulder County (Adams et al. 2018). The manual vetting gave us confidence in Sonobat’s identification with only 4% (1 recording) of the manually vetted recordings that could not be confirmed to species.

### Spatial and temporal patterns

All sites had at least one detection across the active season, but only 50% of sites had detections during all sampling periods (Fig. 2. B-D). Significant differences in mean PPN across sampling periods for five of the 12 sites were detected (Table 1). Of the five sites that did significantly vary, tri-colored bat activity increased for three of the sites throughout the survey period and decreased for the other two.

**Table 1.**
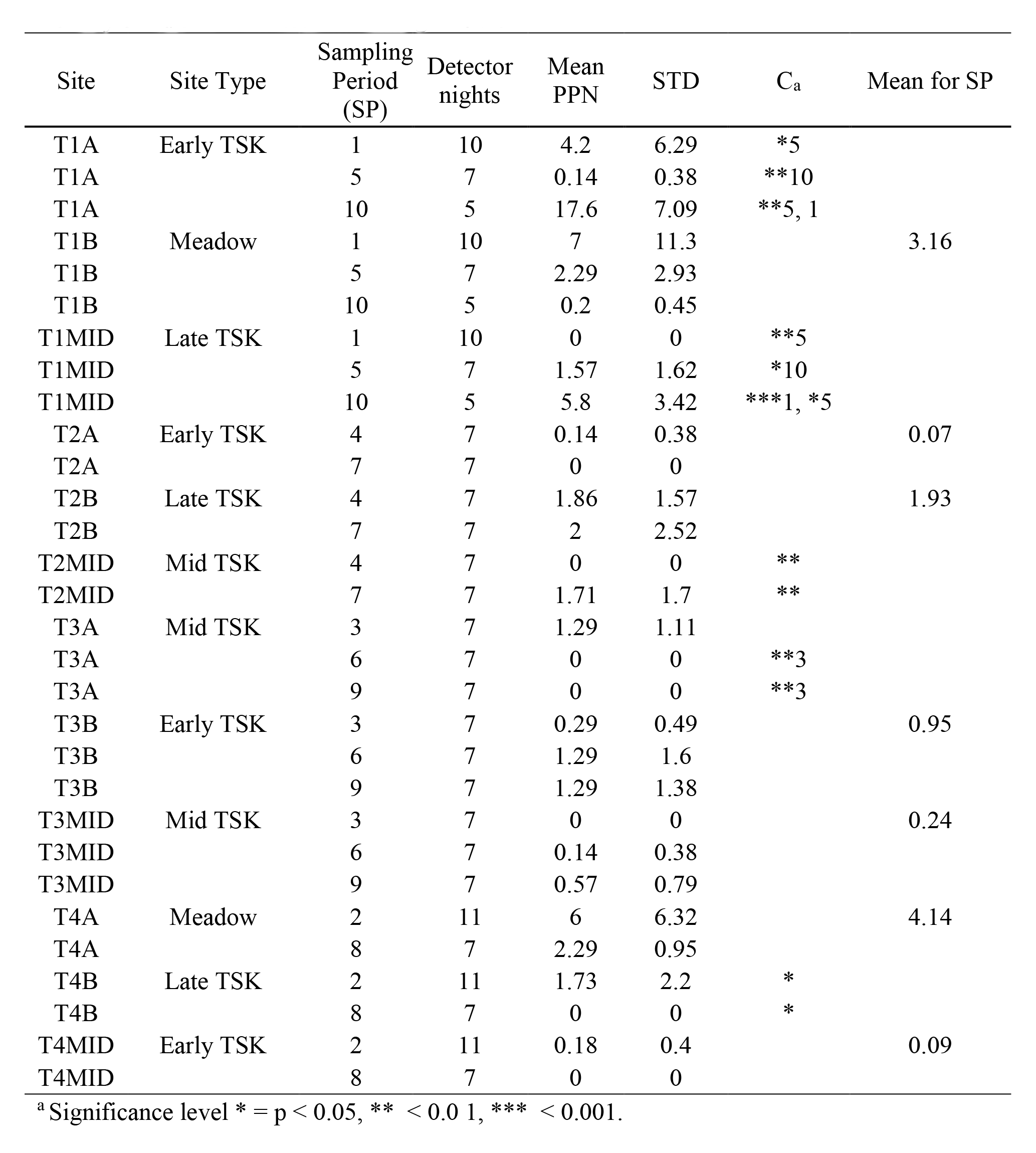
Mean passes per night (PPN) was calculated by summing all tri-colored bat detections for each sampling period (SP) divided by the number of survey nights. STD = standard deviation. Column C indicates which sampling periods have significantly different mean PPN for that site. Note that low SP numbers represent surveys that occurred earlier in the season. Mean PPN for SP was only calculated for sites without significant differences.

### Habitat features used for predictive modelling of bat activity

The NPMR model has a significant fit (xR^2^ = 0.389, p = 0.048), and three variables contributed to model performance. Approximately 39% of variation in tri-colored bat detections were explained by the maximum TSK (sensitivity = 0.048), mean volume of coarse woody debris (CWD) (0.0365), and day of the year (0.49) (Table 2). Day of the year was the most influential variable with a tolerance (standard deviation of the kernel function) of 3.7. Maximum TSK was the second most influential variable with a tolerance of 0.9, and mean volume of coarse woody debris had a tolerance of 0.219. Our model predicted that day of the year had a multimodal relationship with tri-colored bat activity, which is suggestive of interactive effects with other predictors (Fig. 3). Tri-colored bats had higher activity rates at sites with low volumes of CWD during the first two sampling periods but had higher mean PPN in sites with greater volumes of CWD during the final sampling period (Fig. 4A). Tri-colored bats were also predicted to be active in sites with a low maximum TSK throughout the survey period and in sites with a wide range of volumes of CWD (Fig. 4B-C). These results indicated bats may be using a mixture of meadows and early TSK sites which both have zero or low maximum TSK values. We then examined the habitat-specific model estimated activity patterns which predicted that tri-colored bats used a variety of habitat types throughout the survey period (Fig. 5). Meadow sites had the highest estimated mean PPN during sampling periods 1 and 2, but tri-colored bats were predicted to be more active in beetle-killed forests, including both early TSK and late TSK sites, than in meadows during the final sampling period (Fig. 5).

**Table 2.** Nonparametric multiplicative regression model results for predicting tri-colored bat activity in bark beetle-affected forests of Northern Colorado.

| Cross-validated<br>R<sup>2</sup> | N*<br>size<sub>a</sub> | Predictor<br>1 | T1<br>b | S1<sub>c</sub> | Predictor<br>2 | T2<sub>b</sub> | S2<sub>c</sub> | Predictor<br>3 | T3<sub>b</sub> | S3<sub>c</sub> |
| --- | --- | --- | --- | --- | --- | --- | --- | --- | --- | --- |
| 0.388 | 3.01 | Max.TSK | 0.9 | 0.048 | Coarse<br>woody<br>debris<br>(m<sup>3</sup>) | 0.219 | 0.037 | Day of<br>year | 3.7 | 0.49 |

<sup>a</sup> N\* refers to the average data neighborhood size.
<sup>b</sup> T refers to the tolerance (SD of the kernel function) for the preceding predictor.
<sup>c</sup> S refers to the sensitivity for the preceding predictor.

## DISCUSSION

Herein we quantified the relative activity rates of tri-colored bats using higher elevation subalpine forests and meadows, and how tri-colored bats are associated with habitat and landscape variables in areas undergoing secondary succession after severe beetle kill disturbances. We found that tri-colored bat activity is highly variable at these high elevation sites which confirmed prediction 1. We also found that the habitat characteristics associated with activity also varied which confirmed prediction 2. We acknowledge that there is some uncertainty involving confirmation of species identities using acoustic identification. We do, however, feel there is a high degree of certainty in our species identification due to the unique characteristics of tri-colored bat call structure (hockey-stick shaped, ≥40 minimum low frequency, ≤ 70 maximum frequency, etc.), the relatively low overlap in call characteristics with other species in our area, and our ability to compare calls with reference calls gathered in nearby Boulder County by Adams et al. (2018). Although netting individuals would be ideal, the current population density of this species at our high elevation field sites is very low and therefore the probability of capture is equally very low. In addition, the recent discovery of Pd, the fungal agent that causes WNS, in Larimer County has reduced permitting by Colorado Parks and Wildlife for the foreseeable future.

Roosevelt National Forest, where this study took place, has undergone severe bark beetle disturbances in recent years (Klutsch et al. 2009), and our sites included plots affected at various levels (Supplementary Data SD1). Beetle kill disturbances directly and indirectly change habitat features, some of which our models found to be significant predictors of tri-colored bat activity. Our results indicate tri-colored bats were using meadows and forest stands in early TSK stages throughout the active season with highest activity in these habitats in both June and August. Carter et al. (1999) found similar habitat use patterns in eastern deciduous forests, suggesting that this species would avoid the more cluttered vegetation associated with mid TSK stages at our study area. Similarly, Loeb and O’Keefe (2006) found that tri-colored bats were 8.75 times more likely to be found in open patches than in dense forests and they were more likely to be found in early successional stands than mid-successional or mature forest stands with activity lowest in the latter.

We found that habitat features associated with tri-colored bat activity varied depending on time of year with the middle of the survey period (most of July) having the lowest detections of tri-colored bats. However, activity rose sharply towards the end of the survey period, and our model predicted that tri-colored bats use a greater variety of habitat features during this period. Others have also found that tri-colored bats have a highly seasonal use of forest patches and gaps (Loeb & O’Keefe 2006, Braun De Torrez, Ober, & McCleery 2018). Importantly, although tri-colored bats are typically associated with open habitat patches, forest-clearing disturbances appear to decrease activity rates of tri-colored bats, including significant clearing for agriculture or development (Farrow & Broders 2011). These studies, in addition to our own, highlight that tri-colored bats are associated with a mosaic of closed and open landscape features.

It is likely that tri-colored bats are keying into specific landscape features to fit their dietary and changing physiological needs throughout the active season. Reproductive female tri-colored bats have been observed to travel up to 4.3 km from roosting areas to foraging grounds (Veilleux et al. 2003) and use different habitat features, such as various roost types, throughout the summer months (Veilleux et al. 2004). In our study area, tri-colored bat activity associations with forest secondary successional stages may be driven by prey availability. We found that tri-colored bats were active in a wide range of habitats (Fig. 6), but activity was predicted to be highest in sites with high volumes of coarse woody debris at the end of the survey period (sampling period 10, Fig. 5). Coarse woody debris support high diversity and abundance of Hymenopteran, Coleopteran, and Dipteran insects; all of which are important prey for tri-colored bats (Riffell et al. 2011, Feldhamer et al. 2009, Koenigs et al. 2002). However, unknown roosting habits limit our ability to better understand this species’ ecology in novel habitats within its expanded range in Colorado. Tri-colored bats have been documented roosting in dead pine needle clusters in Arkansas (Perry and Thill 2007) and thus might be using beetle-killed pine needles in forest gaps in our study area. Improved technology for acoustic identification has increased the use of acoustic surveys by natural resource managers to identify the presence of tri-colored bats within their newly expanded range (White, Lemen, & Freeman 2016), and although Neaubaum and Agaard (2022) suggest the underuse of subalpine habitats by bats, it is unclear if these results are simply due to a lack of sampling effort. Follow up studies should seek to document the summer roosting ecology of tri-colored bats, and other Rocky Mountain West bat species living in higher elevation forests recovering from large-scale disturbance events.

In conclusion, we showed that tri-colored bats have a sustained but variable presence at relatively high elevation (∼2700m) subalpine habitats. Additionally, the results of this study include descriptions of this species’ expanded niche breadth to include seasonal use of bark beetle killed lodgepole forests in various stages of recovery. Although numbers of echolocation passes for this species are modest at higher elevations, we predict populations will increase over time. Adams et al. (2018) recorded 70.7 mean PPN over ten days or ∼ 7 passes per hour (PPH) in late August along a linear flyway within South Boulder Creek riparian habitat in eastern Boulder County. This is the only published acoustic data from lower elevations in Colorado. Before the onset of WNS, an acoustic survey using ANABAT detectors in Jefferson County and Lewis County, New York in uncluttered riparian habitats indicated about 0.5 PPH (Ford et al. 2011). Given that the average time between sunset and sunrise during our survey period is 9.5 hours, our most consistently active sites (T1B and T4B) experienced comparable activity rates of 0.33 PPH and 0.44 PPH throughout the survey period.

If tri-colored bats can escape WNS by westward expansion and find refuge in novel western habitats where WNS has not become endemic, perhaps recovery of this species will occur. We anticipate that the US Fish and Wildlife Service will list this species as endangered (USFWS 2017) with Federal protections across its range, thereby making Colorado a potential forefront in this species recovery. Unknown habitat requirements for tri-colored bats within its expanded range will hinder conservation efforts for the species. Future studies should seek to investigate the effect of beetle-killed snag removal on tri-colored bat activity in these habitats and determine roost preferences of this species within its expanded range. Additionally, as climate change has led to severe disturbance events in the Rocky Mountain West, future studies should seek to determine how microhabitat characteristics may influence bat use of subalpine habitats as it is unclear how forest bat species will respond to these altered habitats under climate warming scenarios (Hayes & Adams 2017). Through acoustic surveys, we document tri-colored bat activity in novel habitat types in the Rocky Mountain West (disturbed subalpine forests) that will help with species management and protections for westernmost populations in Colorado.

## ACKNOWLEDGEMENTS

This research was financially supported by the University of Northern Colorado. The authors would like to thank Tara Hobbs and Donovan Barratt for their assistance with data collection and two anonymous reviewers for their time and help. U.S. Forest Service research permit #CAN829.

## AUTHOR CONTRIBUTIONS

ABZ and RA conceived the ideas and designed the methodology; ABZ and AC collected the data; ABZ analyzed the data, ABZ, RA, and AC led the writing of the manuscript. All authors contributed critically to the drafts and gave final approval for publication.

## SUPPLEMENTARY DATA

**Supplementary Data SD1. —** Mean habitat and environmental characteristics collected for sites from vegetation surveys and used as model predictors.

**Supplementary Figure Fig. 1 —** A representative sonogram of a tri-colored bat echolocation pass captured in foothills habitats in Boulder County, Colorado (Adams et al. 2018).

---

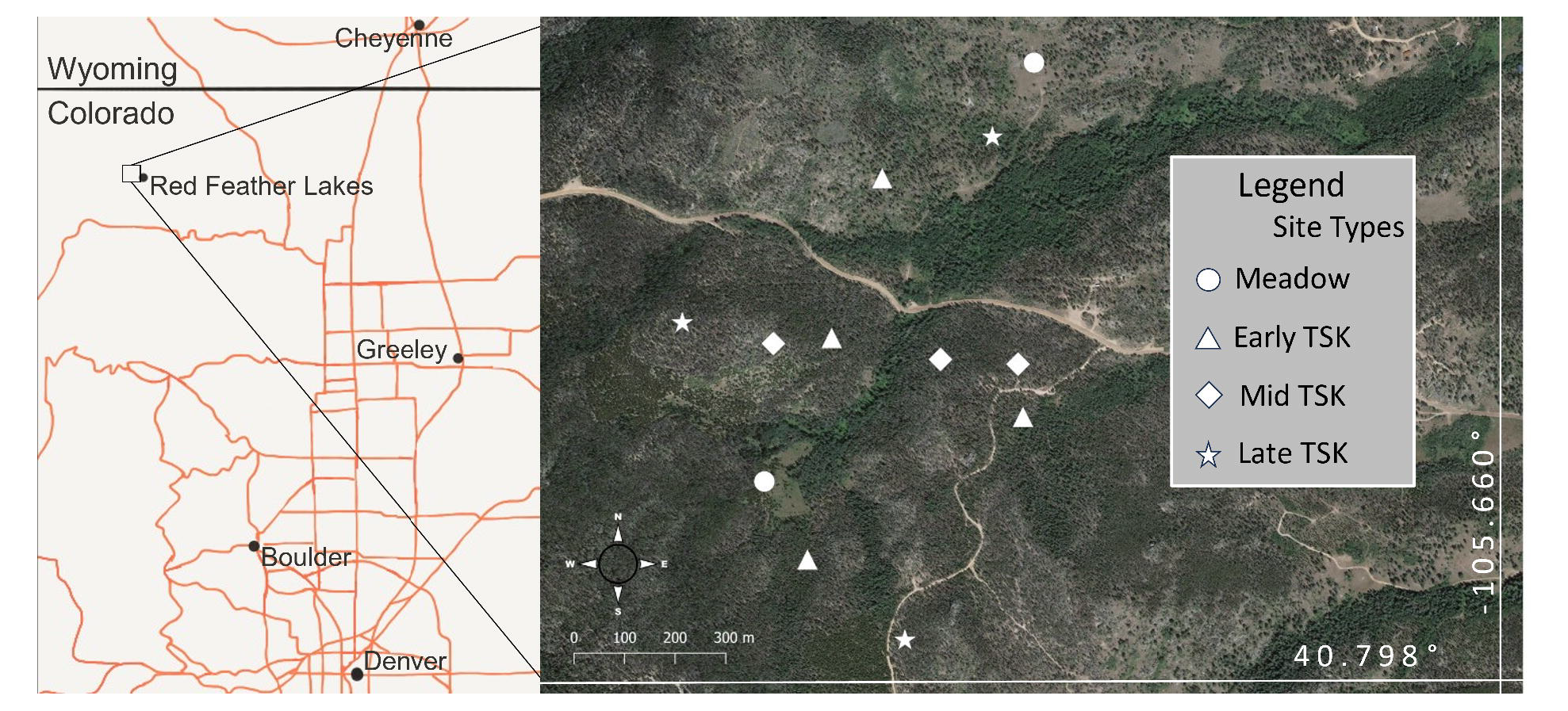

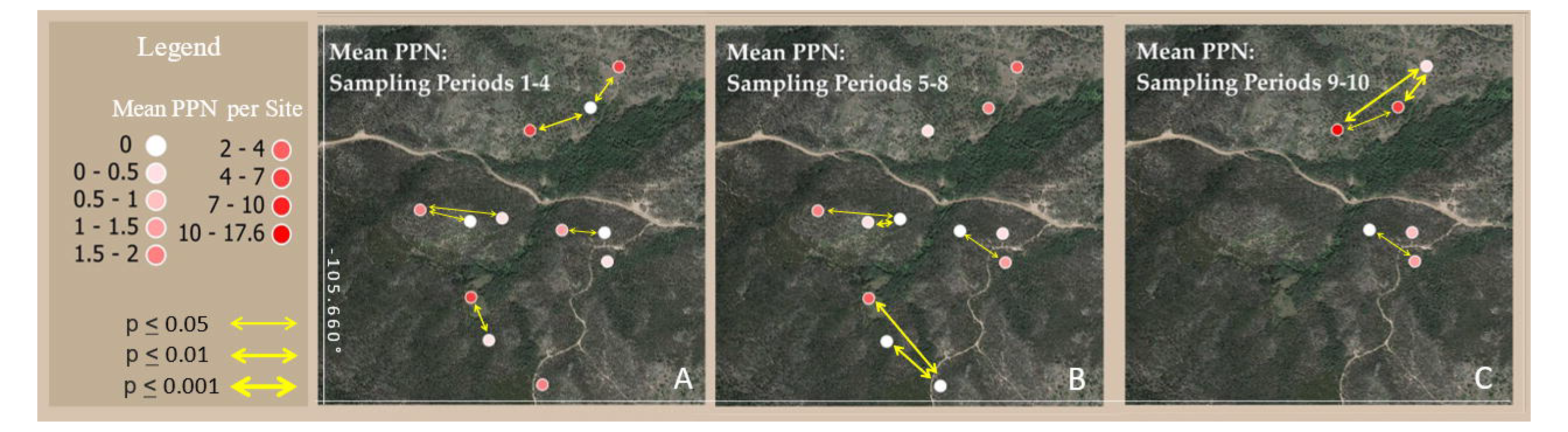

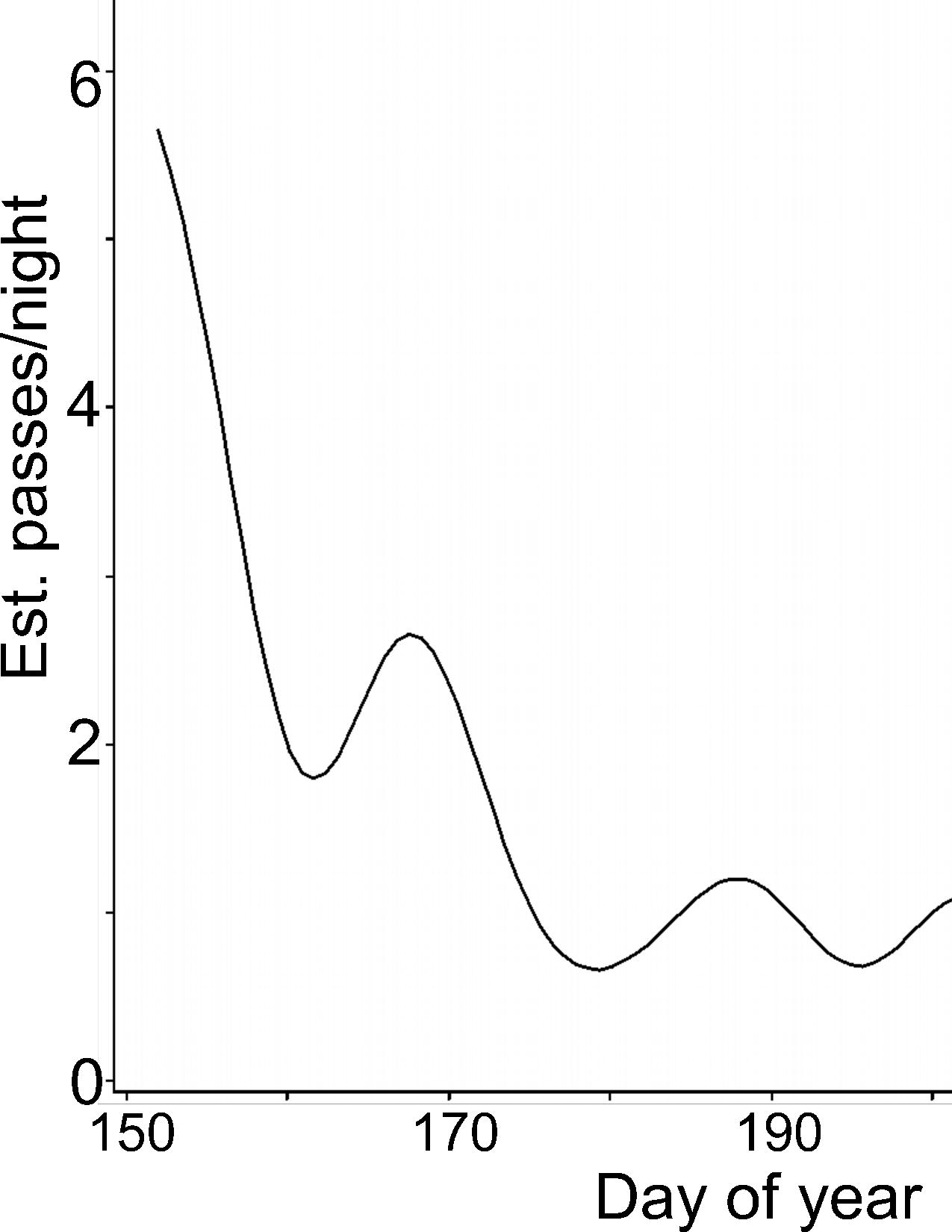

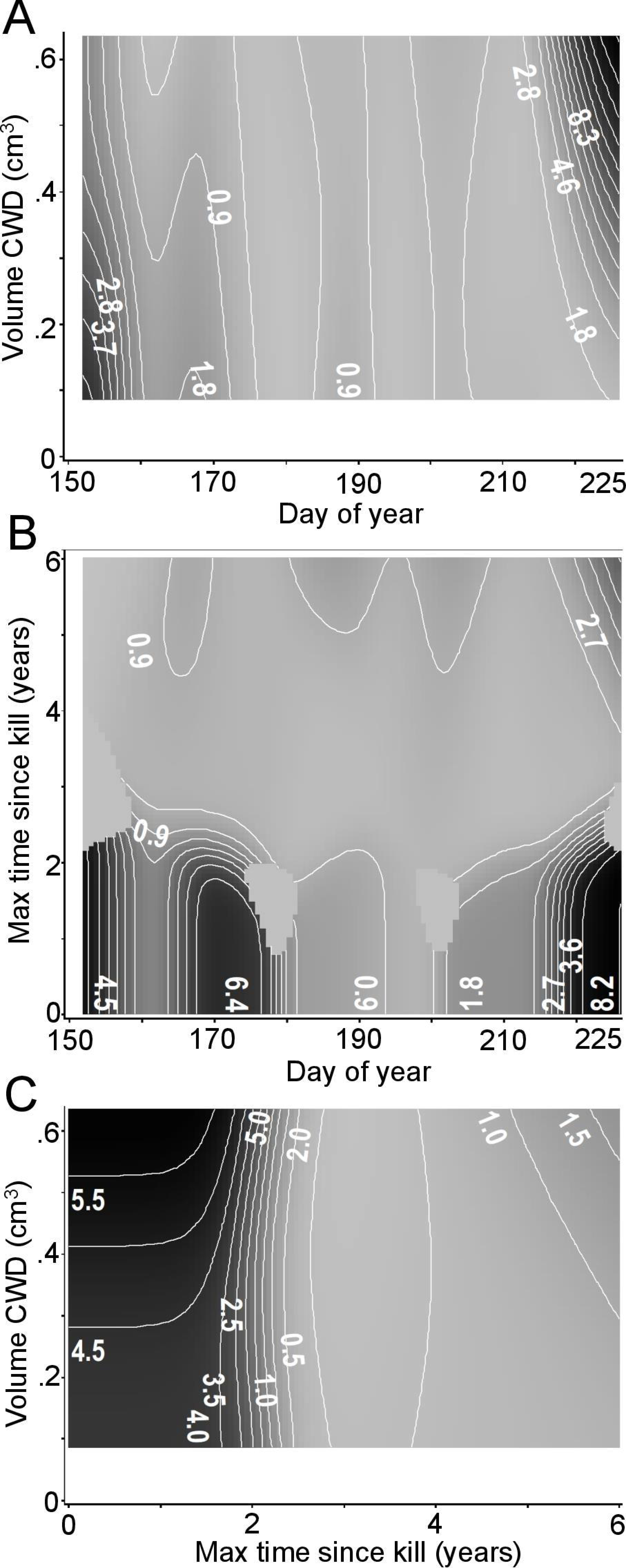

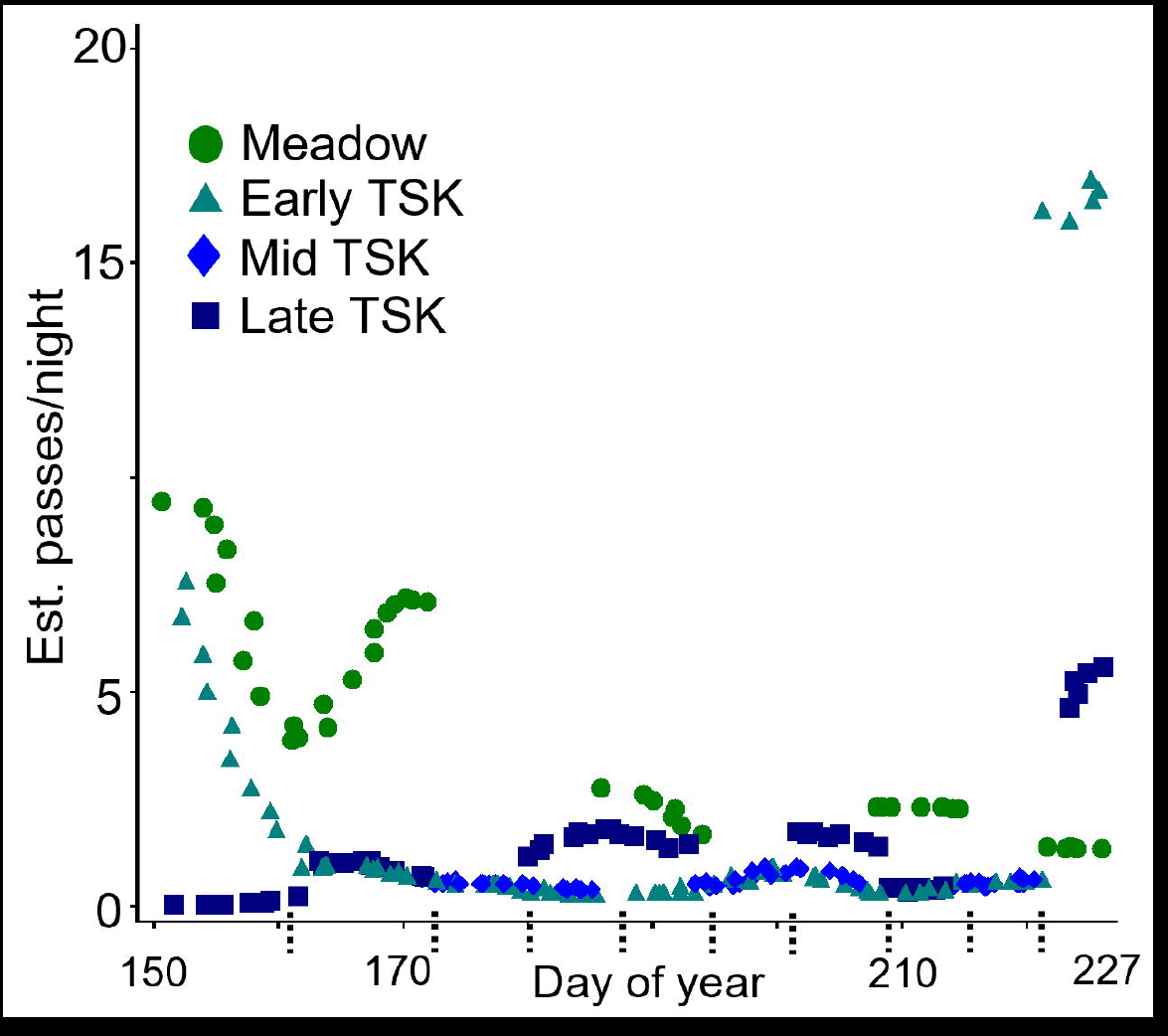

