## Supplementary Data SD1 for "Forest characteristics predict tri-colored bat activity within novel Colorado habitats"

**Supplementary Data SD1. —** Mean habitat and environmental characteristics collected for sites from vegetation surveys.

| **Site** | T1A | T1B | T1MID | T2A | T2B | T2MID | T3A | T3B | T3MID | T4A | T4B | T4MID |
| --- | --- | --- | --- | --- | --- | --- | --- | --- | --- | --- | --- | --- |
| Elevation (m) | 2749 | 2711 | 2705 | 2773 | 2805 | 2799 | 2755 | 2742 | 2743 | 2766 | 2827 | 2790 |
| Aspect, transformed | 0.95 | 0.22 | 0.47 | 0.45 | 0.37 | 0.87 | 0.47 | 0.53 | 0.07 | 0.51 | 0.71 | 0.47 |
| Slope (%) | 26 | 8 | 11 | 11 | 12 | 15 | 25 | 17 | 19 | 11 | 16 | 25 |
| Incident radiation | 0.94 | 0.71 | 0.74 | 0.74 | 0.72 | 0.85 | 0.70 | 0.74 | 0.61 | 0.75 | 0.81 | 0.70 |
| **Stand characteristics** | | | |  |  |  |  |  |  |  |  |  |
| Overstory height (m) | 6.57 | 5.28 | 7.12 | 12.21 | 5.74 | 8.99 | 7.99 | 8.48 | 7.96 | 5.49 | 7.41 | 13.57 |
| Canopy closure | 8.50 | 5.06 | 5.00 | 3.13 | 0.25 | 4.25 | 4.88 | 5.16 | 3.00 | 0.19 | 2.81 | 4.44 |
| Flight corridor width | 5.75 | 29.15 | 1.94 | 3.04 | 6.26 | 3.82 | 3.68 | 5.21 | 4.74 | 17.80 | 2.45 | 4.35 |
| Basal area | 0.21 | 0.30 | 0.24 | 0.18 | 0.23 | 0.18 | 0.19 | 0.17 | 0.22 | 0.05 | 0.28 | 0.27 |
| Beetle kill (%) | 50 | 0 | 56.3 | 77.5 | 63.9 | 60.4 | 65.7 | 52.1 | 56.8 | 0 | 50.4 | 27.0 |
| Average Time Since Kill (TSK) | 0 | 0 | 6.0 | 4.0 | 5.1 | 4.1 | 4.0 | 4.3 | 4.3 | 0 | 5.0 | 4.3 |
| Maximum TSK | 0 | 0 | 6.0 | 4.0 | 6.0 | 4.5 | 4.5 | 5.0 | 5.0 | 0 | 5.0 | 4.5 |
| Mode TSK | 0 | 0 | 0 | 4.0 | 0 | 2.0 | 4.0 | 0 | 2.0 | 0 | 0 | 0 |
| **Understory characteristics** | | | |  |  |  |  |  |  |  |  |  |
| Shrub cover (%) | 18.6 | 5.7 | 16 | 11.7 | 24 | 70.7 | 0 | 5.5 | 6.05 | 0 | 7 | 7.5 |
| Shrub height (m) | 0.09 | 0 | 0.30 | 0.09 | 0.04 | 0.17 | 0 | 0.53 | 0.04 | 0 | 0.02 | 0.05 |
| Sapling cover (%) | 0 | 0 | 21.7 | 0.83 | 7.8 | 3.2 | 6.3 | 17.3 | 2.2 | 1.0 | 2.8 | 0.67 |
| Sapling height (m) | 0 | 0 | 0.68 | 0.02 | 0.36 | 0.12 | 0.23 | 0.60 | 0.09 | 0.05 | 0.14 | 0.03 |
| Grass cover (%) | 2.0 | 23.3 | 9.2 | 0.3 | 4.0 | 7.3 | 7.0 | 1.7 | 2.5 | 31.2 | 7.2 | 0.2 |
| Forb cover (%) | 1.4 | 2.9 | 13.3 | 9.5 | 6.8 | 5.0 | 1.8 | 14.7 | 2.2 | 36.8 | 3.3 | 5.5 |
| Forb height (m) | 0 | 0 | 0 | 0 | 0.09 | 0.28 | 0 | 0.08 | 0.01 | 0 | 0 | 0.01 |
| Moss/lichen cover (%) | 0 | 0 | 0 | 0 | 8.8 | 0 | 0.33 | 0.33 | 9.0 | 0 | 1.0 | 0 |
| Bare ground/ rock cover (%) | 12.0 | 30.0 | 2.2 | 4.2 | 16.5 | 4.0 | 14.5 | 4.5 | 11.7 | 19.3 | 12.8 | 14.2 |
| Medium woody debris (%) | 3.5 | 2.3 | 8.4 | 4.7 | 4.2 | 3.2 | 5.5 | 11.3 | 3.7 | 6.8 | 2.5 | 3.2 |
| Fine woody debris cover (%) | 63.0 | 35.7 | 29.2 | 68.8 | 27.8 | 44.5 | 64.5 | 44.7 | 60.8 | 4.8 | 68.5 | 68.8 |
| **Coarse woody debris volume (cm^3^)** | | | | | |  |  |  |  |  |  |  |
| Total | 0.63 | 0.13 | 0.57 | 0.39 | 0.52 | 0.29 | 0.24 | 0.15 | 0.21 | 0.09 | 0.22 | 0.20 |
| Decay class 1 | 0 | 0 | 0 | 0 | 0 | 0 | 0 | 0 | 0.02 | 0 | 0 | 0 |
| Decay class 2 | 0 | 0 | 0 | 0.22 | 0 | 0 | 0.04 | 0 | 0 | 0 | 0 | 0.07 |
| Decay class 3 | 0 | 0 | 0.04 | 0.09 | 0 | 0.22 | 0.10 | 0 | 0 | 0 | 0 | 0.03 |
| Decay class 4 | 0.44 | 0.08 | 0.20 | 0 | 0 | 0.03 | 0.02 | 0 | 0.08 | 0 | 0 | 0 |
| Decay class 5 | 0.20 | 0.05 | 0.32 | 0.08 | 0.52 | 0.04 | 0.08 | 0.15 | 0.11 | 0.09 | 0.22 | 0.10 |
