## Supplementary figures and images for "Forest characteristics predict tri-colored bat activity within novel Colorado habitats"

### Supplemental Fig.1

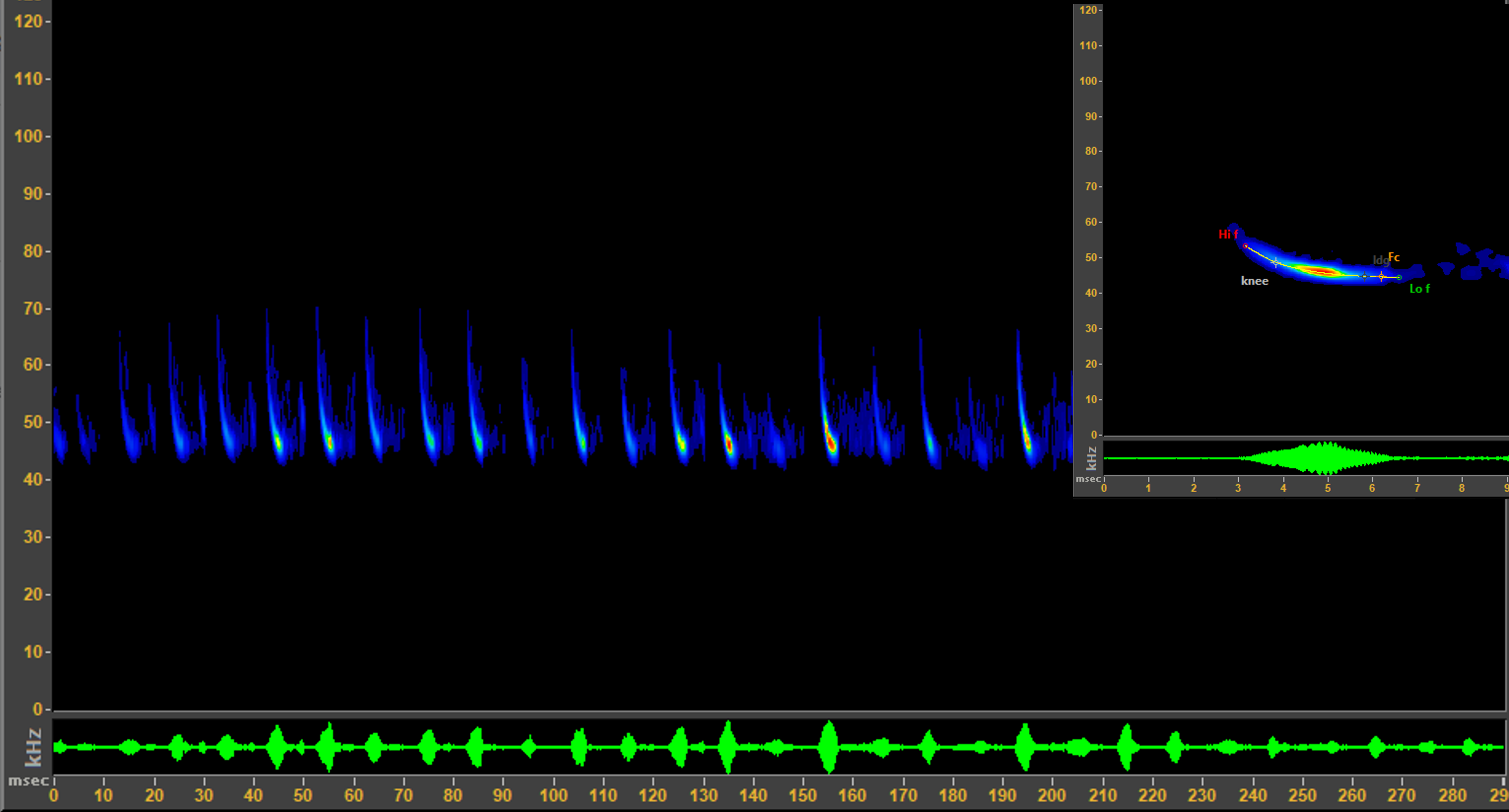
